# Ancient *trans*-acting siRNAs Confer Robustness and Sensitivity onto the Auxin Response

**DOI:** 10.1101/024307

**Authors:** Yevgeniy Plavskin, Akitomo Nagashima, Pierre-François Perroud, Mitsuyasu Hasebe, Ralph S. Quatrano, Gurinder S. Atwal, Marja C.P. Timmermans

**Affiliations:** Watson School of Biological Sciences, Cold Spring Harbor Laboratory, Cold Spring Harbor, NY, USA; National Institute for Basic Biology, Okazaki, Japan; Exploratory Research for Advanced Technology, Japan Science and Technology Agency, Okazaki, Japan; Department of Biology, Washington University, St. Louis, MO, USA; Center for Plant Molecular Biology, University of Tübingen, Tübingen, Germany

**Author notes:** Author of Correspondence: Marja Timmermans, 1 Bungtown Road, Cold Spring Harbor, NY 11724. Present address: Department of Biology, NYU Center for Genomics and Systems Biology, New York, NY, USA. Present address: Department of Cell Biology II, Philipps-Universität Marburg, Marburg, Germany.

**Keywords:** small RNA, evolution, gene regulatory network, auxin, Physcomitrella

## Abstract

Novel developmental programs often evolve via cooption of existing genetic networks. To understand this process, we explored cooption of the *TAS3* tasiRNA pathway in the moss *Physcomitrella patens*. We find an ancestral function for this repeatedly redeployed pathway in the spatial regulation of a conserved set of Auxin Response Factors. In moss, this results in stochastic patterning of the filamentous protonemal tissue. Through modeling and experimentation, we demonstrate that tasiRNA regulation confers sensitivity and robustness onto the auxin response. Increased auxin sensitivity parallels increased developmental sensitivity to nitrogen, a key environmental signal. We propose that the properties lent to the auxin response network, along with the ability to stochastically modulate development in response to environmental cues, have contributed to the tasiRNA-ARF module’s repeated cooption during evolution. The signaling properties of a genetic network, and not just its developmental output, are thus critical to understanding the evolution of multicellular forms.

## INTRODUCTION

The evolution of novel forms is frequently driven by the cooption of existing developmental gene regulatory networks (GRNs) (Erwin and Davidson, 2009). Certain networks appear especially prone to such evolutionary repurposing, leading to the regulation of multiple diverse developmental processes by a single conserved GRN (Carroll et al., 2004a; Plavskin and Timmermans, 2012). The properties that favor the recurring cooption of select GRNs remain largely unknown. One possibility is that frequently coopted GRNs regulate defined cellular processes, allowing the evolutionary redeployment of such ‘differentiation modules’ in a new context (Erwin and Davidson, 2009). On the other hand, it has been postulated that small network motifs may have been repeatedly reutilized by evolution to regulate diverse processes because of the signaling properties they confer (Milo et al., 2002).

The mechanism of network cooption is better understood, with changes impacting the expression of central GRN components frequently driving the evolution of developmental novelties (Carroll et al., 2004b). Small regulatory RNAs, which play a key role in gene regulation during development in both plants and animals, are one mechanism for driving such evolutionary change (Plavskin and Timmermans, 2012). In addition to regulating the level and spatiotemporal expression pattern of their targets, small RNAs are thought to reduce the inherent noisiness of transcription or, as an outcome of mobility, give rise to sharpened gene expression boundaries (Chitwood et al., 2009; Levine et al., 2007; Schmiedel et al., 2015; Skopelitis et al., 2012). These properties may lend robustness to small RNA-regulated networks, perhaps influencing GRN cooptability. An understanding of how the developmental roles of small RNAs and their targets change over the course of evolution may therefore lead to important insights regarding the properties of GRNs that promote their repeated evolutionary cooption.

Plants have undergone tremendous diversification since their colonization of land ~450 million years ago. Some notable innovations include the formation of a lignified vasculature, a sporophyte-dominant life cycle, layered meristems, leaves, flowers, fruits, and seed. While these changes in body plan occurred in parallel with repeated losses and gains of developmentally important small RNAs, select small RNA families and their targets have remained conserved since the most recent common ancestor of all land plants (Axtell et al., 2007; Cuperus et al., 2011). The *TAS3* trans-acting short interfering RNA (tasiRNA) pathway is in this regard of special interest: while conserved throughout land plant evolution, its contributions to development vary extensively even among flowering plants (Plavskin and Timmermans, 2012). *TAS3* tasiRNA biogenesis is triggered in response to the miR390-directed cleavage of a set of non-coding *TAS3* transcripts, which causes conversion of these precursors into long double-stranded RNAs (dsRNAs) by RNA-DEPENDENT RNA POLYMERASE 6 (RDR6) and SUPPRESSOR OF GENE SILENCING 3 (SGS3). The dsRNA intermediates are subsequently processed by DICER-LIKE 4 (DCL4) into 21-nt tasiRNAs that are phased to yield discrete small RNA species. A subset of these tasiRNAs are biologically active and, like miRNAs, act at the post-transcriptional level to regulate expression of specific gene targets (see Plavskin and Timmermans, 2012).

Although related tasiRNA biogenesis pathways initiated by other miRNAs exist (Fei et al., 2013), only the *TAS3* tasiRNA pathway is conserved across land plant evolution (Axtell et al., 2007). Furthermore, only the *TAS3* tasiRNAs have demonstrated roles in development. The TAS3-derived tasiARFs, which regulate the expression of AUXIN RESPONSE FACTORS within the *ARF3* and *ARF4* clade, function in adaxial-abaxial (top-bottom) leaf polarity in a number of flowering plant species, including *Arabidopsis*, tomato, rice, and maize (Chitwood et al., 2009; Nagasaki et al., 2007; Nogueira et al., 2007; Yifhar et al., 2012). In addition, these tasiRNAs contribute to shoot meristem maintenance in monocots (Dotto et al., 2014; Nagasaki et al., 2007), heteroblasty and lateral root outgrowth in *Arabidopsis* (Hunter et al., 2006; Marin et al., 2010; Yoon et al., 2010), and leaf complexity in *Lotus* and *Medicago* (Yan et al., 2010; Zhou et al., 2013). Thus, even within the flowering plant lineage, the *TAS3* tasiRNA pathway has been coopted for the regulation of diverse developmental processes.

Moreover, the organs whose development is regulated by *TAS3* tasiRNAs in flowering plants-leaves, roots, and layered shoot meristems - evolved long after the pathway itself did (Axtell and Bartel, 2005). The genome of the moss *Physcomitrella patens*, which last shared a common ancestor with flowering plants ~450 million years ago, includes six *TAS3* loci (Arif et al., 2012; Axtell et al., 2007). As in flowering plants, a subset of TAS3-derived tasiRNAs in moss target transcripts of ARF transcription factors. This suggests an additional, yet unknown role of *TAS3-*derived tasiRNAs in ancient land plant development. Studying the role of these small RNAs outside the flowering plant lineage thus presents a unique opportunity to shed light on potential ancestral functions of this pathway, and to explore the properties that have led to this pathway’s repeated cooption over the course of plant evolution.

To this end, we generated mutants in *Physcomitrella* that lack SGS3 activity and are impaired in tasiRNA biogenesis. The *Ppsgs3* mutants exhibit defects in gametophore formation, protonemal branching, and differentiation of specialized caulonemal filaments, which result from the upregulation of a conserved set of repressor *ARF* genes at the edge of the developing protonema. We show that tasiRNAs act to generate differential levels of *ARF* expression at the protonemal edge, leading to a stochastic pattern of protonemal cell fate determination. Through a combination of computational modeling and experimentation, we further demonstrate that tasiRNA regulation confers sensitivity and robustness onto the auxin response. Finally, in line with this pathway’s role in tuning the auxin sensitivity of cells at the plant’s growing edge, we find that *Physcomitrella* plants defective in tasiRNA biogenesis display decreased developmental sensitivity to a key environmental signal. We propose that the properties lent to the auxin response gene regulatory network by tasiRNAs, along with their ability to stochastically modulate development in response to environmental cues, have contributed to the repeated cooption of the tasiRNA-ARF module over the course of plant evolution.

## RESULTS

### Disruption of tasiRNA biogenesis in *Physcomitrella* impairs protonemal development

*Physcomitrella* homologs of *RDR6* and *DCL4* with a conserved role in tasiRNA biogenesis have been described previously (Arif et al., 2012; Cho et al., 2008; Talmor-Neiman et al., 2006). However, the contributions of tasiRNAs to moss development remain unresolved, in part because mutations perturbing these biogenesis components give rise to disparate phenotypes. To gain insight into the developmental role of the *TAS3* tasiRNA pathway outside the flowering plant lineage, we used targeted recombination to disrupt the single *Physcomitrella* homolog of *SGS3, PpSGS3* (Figure S1A-E). Consistent with a conserved role of PpSGS3 in tasiRNA biogenesis, levels of these small RNAs are drastically reduced in *Ppsgs3* (Figure 1A). The *Physcomitrella* genome includes six *TAS3* loci, *PpTAS3a-f*. These, in contrast to their counterparts in flowering plants, yield two sets of biologically active small RNAs: tasiARF, which target transcripts of four *ARF* genes, *PpARFb1-4*, and tasiAP2, which regulate expression of members in the APETALA2 (AP2) transcription factor family (Arif et al., 2012; Axtell et al., 2007; Talmor-Neiman et al., 2006). Levels of both biologically active tasiRNAs are drastically reduced in *Ppsgs3*, whereas levels of miR390, which functions upstream of SGS3 in tasiRNA biogenesis, remain unaffected **(Figure 1A)**. In addition, the tasiRNA-guided cleavage of target transcripts is specifically affected in *Ppsgs3*. Expression of the *PpARFb* genes is regulated by tasiARF as well as the miRNA miR1219 (Axtell et al., 2007). 5’ RACE analysis revealed that while the miR1219-guided cleavage of *PpARFb1* and *PpARFb4* transcripts is not noticeably affected, the accumulation of tasiARF-directed cleavage products for both transcripts is strongly reduced in *Ppsgs3* plants (Figure S2A). Together with the striking reduction in tasiRNA levels in *Ppsgs3*, this indicates that a role of SGS3 in tasiRNA biogenesis is conserved across land plants.

**Figure 1.**
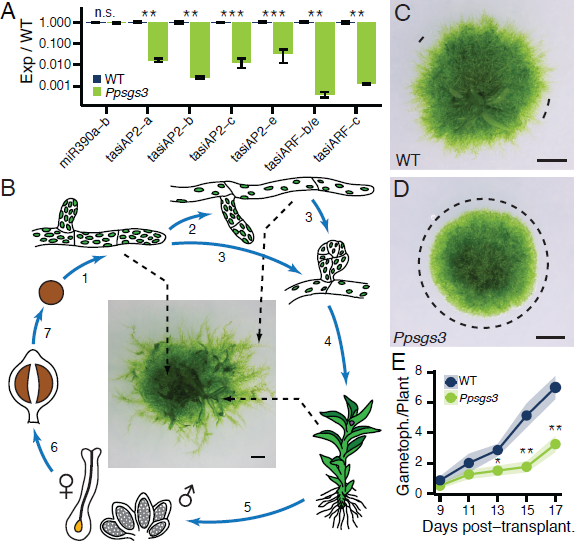
*Ppsgs3* mutants defective in tasiRNA biogenesis exhibit defects in gametophyte development. (A) Levels (mean ± SE, n ≥3) of miR390 and TAS3-derived tasiRNAs in *Ppsgs3* normalized to wild type; **P < 0.01, ***P < 0.001, Student’s *t* test. (B) Overview of the moss life cycle. 1) A germinating haploid spore gives rise to chloronemal filaments that by tip growth and branching form a dense network. 2) After ∼2 weeks, random tip cells differentiate to form long caulonemal filaments. 3) Both filaments also form modified side branches, 4) which grow into leafy gametophores. 5) These carry the sexual organs, and 6) upon fertilization the diploid sporophyte that 7) produces new spores via meiosis. (C-D) Caulonema occurrence on 15 day-old wild type (C) and *Ppsgs3* (D) plants. Dotted lines, areas at the protonemal edge lacking caulonema. Scalebar, 1mm. (E) Gametophore numbers (mean ± SE, n ≥10) in *Ppsgs3* compared to wild type. *P < 0.05, **P < 0.01, Student’s *t* test. See also Figures S1 and S2.

Loss of PpSGS3 activity results in defined developmental defects. While *Ppsgs3* plants show no obvious sporophyte defects and produce viable spores (**Figure S2B, C**), protonemal development in the mutant is perturbed. Following germination, wild type moss spores produce filamentous, branching protonema comprised initially of chloroplast-rich chloronema (Reski, 1998; **Figure 1B**). However, as development progresses, or in response to certain environmental cues, dividing chloronemal cells may transition to give rise to the more elongated caulonemal cells. In addition, modified protonemal side branches also give rise to buds, which in turn develop into leafy gametophores. While indistinguishable from wild type early in development, the chloronemal networks of 15-day-old *Ppsgs3* mutants are smaller and denser than those of wild type (**Figure 1C, D**), due in part to a decrease in both chloronemal cell size and branch determinacy (**Figure S2H-L**).

The most striking phenotype of *Ppsgs3* mutants, however, concerns the formation of long caulonemal filaments. Under the growth conditions used, long caulonemal filaments visibly extend from the chloronemal edge of wild type plants after approximately two weeks of development, causing older plants to take on a fuzzy appearance (**Figure 1C**). Although cells with diagonal cross-walls reminiscent of caulonema are occasionally observed in the protonemal network of *Ppsgs3* plants, these mutants either completely lack long caulonemal filaments, or rarely form small isolated patches of such filaments along the otherwise smooth protonemal edge (**Figure 1C, D**). Even after two months of growth, *Ppsgs3* plants rarely form long caulonemal filaments, indicating a suppression of caulonemal filament formation rather than a delay in the chloronema-to-caulonema transition upon loss of SGS3 function.

In addition to these protonemal phenotypes, *Ppsgs3* plants form significantly fewer leafy gametophores than wild type (**Figure 1E**), although gametophore morphology itself is normal (**Figure S2D-G**). This defect is not fully explained by the lack of long caulonemal filaments in *Ppsgs3*, as a significant decrease in gametophore number is detected prior to the normal appearance of these filaments at ∼15 days of growth. Instead, their reduced numbers indicate a role for PpSGS3 in gametophore formation as well as protonemal development. While a reduction in caulonema formation has also been reported for *Pprdr6*, the observed decrease in gametophore number in *Ppsgs3* is at odds with previous data showing that gametophore formation is accelerated in *Pprdr6* compared to wild type (Talmor-Neiman et al., 2006). However, this aspect of the *Pprdr6* phenotype likely reflects its role in 22-24 nt siRNA biogenesis, as a similar defect is seen in *Ppdcl3*, which is required for the production of 22-24 nt siRNAs but not tasiRNAs (Cho et al., 2008).

Taken together, our results indicate that a role for SGS3 in tasiRNA biogenesis is conserved between *Physcomitrella* and flowering plants. PpSGS3 is required for normal gametophyte development, with *Ppsgs3* mutants displaying defects in chloronemal cell size and branch determinacy, caulonemal differentiation, and gametophore formation.

### tasiRNAs affect gametophyte development by regulating expression of *PpARFb* genes

In flowering plants, tasiRNAs exert their effect on development by regulating expression of ARF3 and ARF4 transcription factors (Chitwood et al., 2009; Dotto et al., 2014; Hunter et al., 2006; Yifhar et al., 2012; Zhou et al., 2013). In *Physcomitrella*, TAS3-derived tasiRNAs target *ARF* and AP2 transcription factor transcripts (Axtell et al., 2007; Talmor-Neiman et al., 2006). In addition, an apparent species-specific *TAS* pathway exists in moss. The *Physcomitrella* genome includes *TAS* loci whose transcripts are processed in a miR156- and miR529-dependent manner, referred to as *TAS6*, which yield tasiRNAs targeting a ZF-domain transcription factor (Arif et al., 2012; Cho et al., 2012). To elucidate which tasiRNA targets are responsible for the developmental defects observed in *Ppsgs3*, we compared the expression levels of known tasiRNA targets between wild type and *Ppsgs3* in 15-day-old plants, when all aspects of the *Ppsgs3* phenotype are apparent. Of the four tasiARF-regulated *ARF* genes in *Physcomitrella*, only *PpARFb1, PpARFb2*, and *PpARFb4* are expressed during the gametophyte stage of development. In *Ppsgs3* plants, transcript levels of all three *ARF* genes are upregulated ~2- to 3- fold relative to wild type (**Figure 2A**). In contrast, expression levels of the tasiAP2 and tasiZF targets are not significantly changed between wild type and *Ppsgs3* at this developmental stage (**Figure 2A**), suggesting that the *Ppsgs3* developmental defects result, at least in part, from a failure to correctly regulate the tasiARF targets.

**Figure 2.**
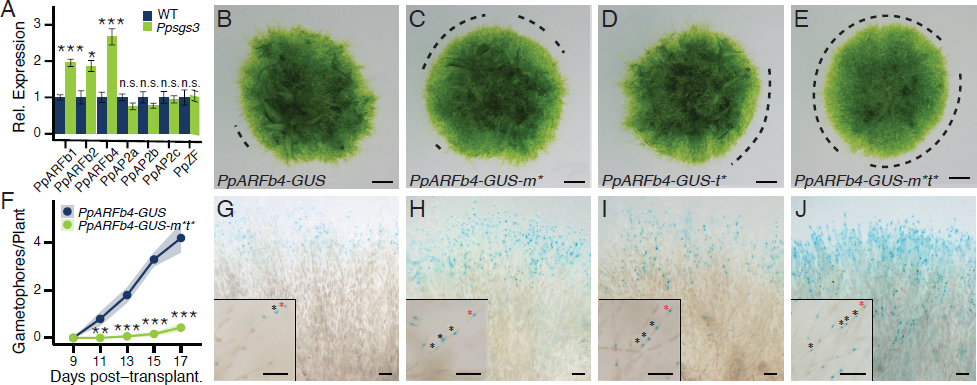
The *Ppsgs3* phenotype reflects misregulation of tasiARF-targeted ARFs at the protonema edge. (A) qRT-PCR values for tasiRNA targets in *Ppsgs3* (mean ± SE, n ≥3) normalized to wild type. *P < 0.05, ***P < 0.001, Student’s *t* test. (B-E) Caulonema occurrence on 15 day-old *PpARFb4-GUS* (B), *PpARFb4-GUS-m\** (C), *PpARFb4-GUS-t\** (D), and *PpARFb4-GUS-m*t\** (E) plants. Dotted lines, protonemal regions lacking caulonema. Scalebar, 1mm. (F) Gametophore numbers (mean ± SE, n $10) on *PpARFb4-GUS* and *PpARFb4-GUS-m*t\** plants. **P < 0.01, ***P < 0.001, Student’s *t* test. (G) *PpARFb4-GUS* activity in 1-3 cells at the tips of a small random (Ljung-Box test, P = 0.19) subset of chloronemal filaments. (H-J) Expanded reporter activity in *PpARFb4-GUS-m\** (H), *PpARFb4-GUS-t\** (I), and *PpARFb4-GUS-m*t\** (J). The latter reflects the broad activity of the *PpARFb4* promoter. Insets show isolated filaments; asterisks, cells expressing PpARFb4-GUS; red asterisks, tip cells. Scalebar, 0.1mm. See also Figure S4.

To investigate this hypothesis, we generated plants that allow the estradiol-inducible expression of HA-tagged, miR1219- and tasiARF-resistant (m*t*) versions of *PpARFb2* and *PpARFb4* (**Figure S3A-C**), which represent the two distinct branches of the *PpARFb* clade (Plavskin and Timmermans, 2012). Even when grown on low concentrations of estradiol, the phenotype of plants overexpressing *PpARFb2-m*t\** or *PpARFb4-m*t\** resembles that of *Ppsgs3*. Such plants form denser protonema with fewer gametophores and suppress the transition from chloronemal to caulonemal development (**Figure 3C-F**). Similar phenotypes are not observed in plants expressing a GUS-GFP fusion protein upon estradiol treatment (**Figure 3A, B**). This indicates that PpARFb2 or PpARFb4 overexpression is sufficient to recapitulate the *Ppsgs3* phenotype, which lends support to the hypothesis that loss of tasiARF activity and the correct regulation of its *PpARFb* targets underlies the developmental defects of *Ppsgs3* mutants.

To substantiate the above findings and to explore the interaction between tasiARFs and miR1219 in the regulation of *PpARFb* expression, we generated a translational fusion of the endogenous *PpARFb4* gene to the GUS reporter, and introduced mutations that prevent targeting of *PpARFb4-GUS* transcripts by miR1219 (m*), tasiARF (t*), or both small RNAs (m*t*) (**Figure S4A-E**). *PpARFb4* was chosen because its expression is especially sensitive to changes in tasiARF regulation (**Figures 2A; S2A**). *PpARFb4-GUS* plants show a subtle decrease in the number of caulonemal filaments but otherwise look phenotypically normal (**Figure 2B**). Likewise, plants expressing the *PpARFb4-GUS-m\** or *PpARFb4-GUS-t\** variants show a variable decrease in the number of long caulonema; however, neither fully recapitulates the *Ppsgs3* phenotype (**Figure 2C, D**). Thus, even though *PpARFb4* transcript levels are increased approximately 3-fold in *Ppsgs3*, loss of tasiARF-mediated regulation of *PpARFb4* alone is insufficient to recapitulate the *Ppsgs3* phenotype. This suggests that altered expression of multiple *PpARFb* targets is needed to condition the developmental defects observed in *Ppsgs3*. Indeed, the phenotypic similarities between *PpARFb2-m*t\** and *PpARFb4-m*t\** overexpressing plants (**Figure 3D, F)** indicates that proteins within this clade have related activities.

**Figure 3.**
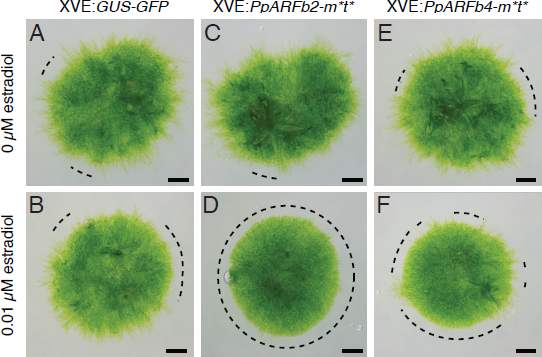
Plants overexpressing *PpARFb2* or *PpARFb4* resemble *Ppsgs3* in phenotype. (A-F) Caulonema occurrence on 15 day-old plants following the estradiol-inducible expression of GUS-GFP (A, B), *PpARFb2-m*t\** (C, D) or *PpARFb4-m*t\** (E, F). Plants grown on 0μM (A, C, E) or 0.01 μM (B, D, F) β-estradiol. Scalebar, 1mm. See also Figure S3.

Importantly, *PpARFb4-GUS-m*t\** plants fail to form long caulonemal filaments (**Figure 2E**), closely mimicking the phenotype observed in *Ppsgs3*. Likewise, *PpARFb4-GUS-m*t\** mutants show a significant and very strong decrease in gametophore number, with the onset of gametophore formation delayed approximately one week relative to plants expressing the small RNA-sensitive *PpARFb4-GUS* fusion (**Figure 2F**). These results indicate that tasiARF and miR1219 coordinately regulate *PpARFb4* expression during gametophyte development. Moreover, the phenotype of *Ppsgs3* mutants, in which expression of three *PpARFb* genes is increased due to loss of tasiARF activity, can be phenocopied by perturbation of tasiARF and miR1219 regulation of *PpARFb4* alone, as well as by the inducible overexpression of a similar mutant version of *PpARFb2* (**Figure 3D**). It follows that tasiARF, in concert with miR1219, regulates the chloronema-to-caulonema transition and gametophore formation in *Physcomitrella* by fine-tuning the combined level of *PpARFb* expression. When *PpARFb* levels overall exceed a certain level, not reached in *PpARFb4-GUS-t\** plants, formation of long caulonemal filaments and gametophores is inhibited.

### tasiARF and miR1219 limit PpARFb4 expression at the protonemal edge

Small RNAs make diverse contributions to plant development (see Skopelitis et al. (2012)). They are thought to refine levels of protein accumulation by dampening the noise in target gene expression, and can limit expression of their developmental targets to defined spatial and/or temporal domains. Considering that the formation of caulonema and leafy gametophores is in part temporally regulated, tasiARF and miR1219 might control the development of these tissues by regulating the temporal window of *PpARFb* expression. To assess this possibility, we monitored their accumulation over 3 weeks of gametophyte growth. While expression levels for tasiARF, miR1219, and miR390 are temporally regulated and increase up to ∼60 fold during that developmental time window (**Figure S4F**), transcript levels for *PpARFb1, PpARFb2*, and *PpARFb4* change comparatively little during that time (**Figure S4G**).

This observation argues against a role for tasiARF and miR1219 solely in the temporal regulation of *PpARFb* genes, and instead suggests that these small RNAs may act to maintain a spatial domain of *PpARFb* expression. To examine this possibility, we compared the pattern of PpARFb4-GUS activity in plants expressing the wild type or small RNA-resistant variants of this reporter. When regulated by tasiARF and miR1219, *PpARFb4-GUS* expression is limited to 1-3 cells nearest the filament tip at the outer edge of the protonema. Expression in these cells is punctate, consistent with PpARFb proteins functioning as nuclear-localized transcription factors (**Figure 2G**). Interestingly, *PpARFb4-GUS* expression is not seen in all chloronemal filaments, but instead appears variable, with a random subset of filaments along the circumference of the plant showing reporter activity (**Figure 2G**). Considering that increased *PpARFb* expression represses caulonemal filament formation, stochasticity in PpARFb4 levels along the protonemal edge may be linked to the sporadic nature of caulonemal filament formation, which initially appears in seemingly random patches at the edge of the protonema (e.g. **Figure 2B**).

Mutation of either the tasiARF or the miR1219 target site results in increased *PpARFb4* expression, with a greater number of chloronemal filaments and a greater number of cells per filament showing reporter activity (**Figure 2H, I**). Consistent with the stronger phenotype of *PpARFb4-GUS-m*t\** plants (**Figure 2E**), reporter activity is expanded even further upon mutation of both small RNA target sites. In these mutants, PpARFb4-GUS expressing cells occur in nearly every chloronemal filament, and expression extends further down the filament than in either the wild type, *PpARFb4-GUS-m\** or *PpARFb4-GUS-t\** lines (**Figure 2J**). In addition, PpARFb4-GUS levels in individual cells appear stronger than in other genotypes. Importantly, the pattern of *PpARFb4* expression in all genotypes does not change substantially from 8 to 15 or 22 days of growth. This finding is consistent with the transcript level analysis (**Figure S4G)** and supports the hypothesis that tasiARF and miR1219 do not establish a temporal pattern of *PpARFb* expression in protonema. Instead, the data shows that tasiARF and miR1219 act coordinately along the chloronemal filament to limit the spatial expression domain of *PpARFb4* to the outer edge of the growing protonema. Considering that *PpARFb1* and *PpARFb2* expression also increases in *Ppsgs3*, it seems likely that tasiARF and miR1219 regulate these targets similarly.

Together, these data provide a basis for the caulonemal defect of *Ppsgs3* mutants, and suggest a mechanism by which tasiARF, in concert with miR1219, regulates the chloronemal-to-caulonemal transition. Caulonemal filaments differentiate from chloronemal tip cells. tasiARF and miR1219 regulate this process by limiting expression of PpARFb transcription factors, which act as repressors of caulonemal differentiation, to the outer edge of growing protonema. The combined activity of these small RNAs appears finely balanced such that it generates a randomly variegated pattern of PpARFb expression at the protonemal edge. In a subset of chloronemal filaments, PpARFb activity is depleted even in the tip cell, allowing their differentiation into caulonema. Upon loss of small RNA regulation, whether by disruption of tasiRNA biogenesis or small RNA complementarity within *PpARFb* transcripts, expression of these ARF proteins persists in a broader domain that encompasses a larger number of chloronemal tip cells, preventing their differentiation into caulonemal filaments. Interestingly, our data suggest that an absence of *PpARFb* expression is not sufficient for caulonema differentiation, as a variegated pattern of PpARFb4-GUS activity at the protonemal edge is also observed in young plants not initiating caulonemal filaments. Rather, it appears that tip cells lacking PpARFb expression are competent to differentiate into caulonema, but only do so upon receipt of a distinct signal.

### tasiRNAs target repressor ARF transcripts to modulate the auxin response

ARF transcription factors, including the tasiARF targets, form part of a highly conserved GRN that regulates the response to the phytohormone auxin. This network integrates auxin perception into development by controlling the transcription of auxin-responsive genes (ARGs). ARF proteins, which have been classified as either “activator” or “repressor” based on their effect on ARG expression, bind the promoters of such genes in an auxin-independent manner. Transcription factor activity of activator ARFs is, however, blocked in the absence of auxin through dimerization with Aux/IAA proteins. The latter are degraded in response to auxin, resulting in the auxin-dependent derepression of ARG expression (Finet and Jaillais, 2012). Interestingly, the Aux/IAA genes are themselves transcribed in response to auxin signaling, forming a negative feedback loop in the auxin response GRN that is conserved between moss and flowering plants (Prigge et al., 2010). The repressor ARFs add additional complexity to this network, with current models suggesting that repressor ARFs compete with activator ARFs for binding to ARG promoters to allow for the differential regulation of the transcriptional auxin response in space and time (Vernoux et al., 2011).

As in flowering plants, auxin regulates a diverse set of developmental processes in moss, including those impacted by the tasiRNA pathway (Prigge et al., 2010). Indeed, both *Ppsgs3* and lines misexpressing the *PpARFb* targets mimic phenotypes resulting from the treatment of *Physcomitrella* with the ‘anti-auxin’ compound p-Chlorophenoxyisobutyric acid (PCIB) (**Figure S5A, B**). The phenotypes of these mutants are also similar to those of classical auxin-insensitive mutants in moss, which form densely packed chloronema, lack caulonemal filaments, and develop fewer or no gametophores (Prigge et al., 2010). These similarities suggest that the PpARFb proteins may act as repressors of the auxin response. This hypothesis is supported by phylogenetic analysis, which places the *Physcomitrella* tasiRNA-targeted *ARF*s sister to the B group of *ARF* genes in flowering plants that includes known repressor ARFs, such as the tasiARF target *ARF3* (Plavskin and Timmermans, 2012). Indeed, consistent with a role of the *Physcomitrella* tasiRNA targets in repressing the auxin response, transcript levels of the early auxin response genes *PpIAA1a* and *PpIAA1b* (Prigge et al., 2010) are reduced in *Ppsgs3* compared to wild type (**Figure S5C**).

The role of PpARFb proteins as repressors of the auxin response provides additional insight into the stochastic nature of caulonema formation. The tasiARF- and miR1219-generated variation in PpARFb expression at the protonemal edge results in variable responsiveness of chloronemal tip cells to auxin, which is known to promote caulonemal fate. Importantly, the phenotypes of tasiRNA biogenesis mutants in flowering plants also result from changes in the spatiotemporal expression of B-group ARF genes (Chitwood et al., 2009; Dotto et al., 2014; Hunter et al., 2006; Yifhar et al., 2012; Zhou et al., 2013). By regulating the pattern and level of repressor ARF accumulation, the tasiRNA pathway may thus act to modulate the auxin response across spatial domains in the plant; this finding sheds light on a potential ancestral function of this pathway.

### Auxin regulation of *PpARFb* genes creates a second negative feedback loop in the auxin response network

Considering the conserved role of the tasiRNA pathway in regulating canonical repressors of the auxin response, an understanding of how tasiRNAs and their targets affect the signaling properties of the ancient auxin response network at the cellular level may elucidate potential reasons for the tasiARF-ARFb module’s repeated evolutionary cooption. A correct model of the architecture of the auxin response GRN is, in this regard, key. An important conserved feature of this network is the negative feedback between Aux/IAA genes and the auxin response (Prigge et al., 2010). In *Arabidopsis*, expression of a subset of repressor ARF genes is also subject to feedback from the auxin response (Marin et al., 2010; Yoon et al., 2010). As this may impact a GRN’s signaling properties, we determined whether auxin signaling affects expression of the *PpARFb* tasiRNA targets in *Physcomitrella* by analyzing *PpARFb1, PpARFb2*, and *PpARFb4* transcript levels in plants grown on media containing 0.1 µM NAA. Transcript levels for all three genes are increased ~2.5-fold in auxin-grown plants (**Figure 4A**). Consistent with this finding, PpARFb4-GUS expression is expanded in plants grown on auxin-supplemented media, with more protonemal filaments and more cells per filament showing reporter activity (**Figure S5F, H**). Increased expression in response to auxin is also observed for the tasiRNA-resistant form of *PpARFb4-GUS* (**Figure S5G, I**), suggesting that auxin promotes *PpARFb* expression at the transcriptional level, rather than exclusively via repression of tasiARF species. Indeed, the small RNAs regulating *PpARFb* expression show a complex response to auxin treatment. Although some tasiARF species are upregulated in plants grown on media supplemented with auxin, other tasiARF species, as well as miR1219, are repressed (**Figure S5J**). These data establish the existence of a negative feedback loop between auxin and repressor *PpARFb* levels that, in addition to the highly conserved negative feedback between auxin signaling and Aux/IAA genes, modulates the *Physcomitrella* auxin response GRN (**Figure 4B**). Interestingly, this network configuration appears to be partially conserved, as the *Arabidopsis* tasiARF target *ARF4* is also upregulated in response to auxin (Marin et al., 2010; Yoon et al., 2010).

**Figure 4.**
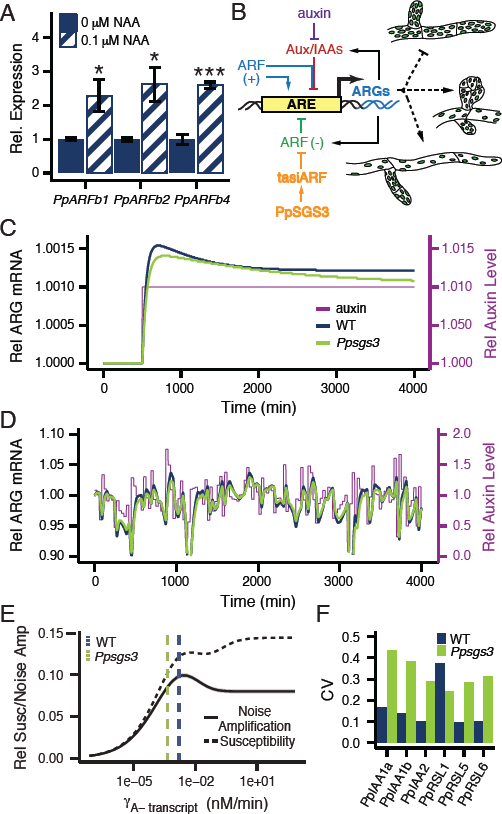
Noise-buffering properties of the auxin response GRN. (A) *PpARFb* transcript levels (mean ± SE, n ≥3) in 15 day-old plants grown on media with 0.1 versus 0μM NAA. *P <0.05, ***P <0.001, Student’s *t* test. (B) Schematic of the *Physcomitrella* auxin response network. Auxin promotes degradation of Aux/IAA proteins, which repress ARG expression through interaction with activator ARFs (ARF (+)). ARG transcription is also repressed by repressor ARFs (ARF (-)), whose levels are modulated by tasiARF activity. ARG transcription only occurs when activator ARFs occupy the Auxin-Responsive Element (ARE) in the promoter. The network contains two negative feedbacks, with both Aux/IAAs and repressor ARFs upregulated in response to auxin. The auxin response represses protonemal branching and promotes caulonema and bud formation. (C) Simulation of ARG expression in response to a prolonged, steady auxin signaling input under both ‘wild type’ and *’Ppsgs3’* conditions plotted relative to the pre-signal state. (D) Simulation of ARG expression in response to a noisy auxin signaling input, plotted relative to steady-state ARG expression levels in ‘wild type’ and *’Ppsgs3’*. Auxin level fluctuations are shown relative to the mean auxin level. (E) Effect of increasing ARF(-) transcript degradation rate, as a measure for small RNA regulation, on noise amplification (mean ± SE) and susceptibility. γ_A-transcript_ values for ‘wild type’ and *’Ppsgs3’* used in (C) and (D) are marked. (F) The expression level CV for six ARGs in wild type and *Ppsgs3*. See also Figure S5.

Repressive feedback from the tasiARF-targeted ARFs onto the auxin response may impart important properties to the auxin response GRN in *Physcomitrella*. Negative autoregulatory circuits are central in allowing cells to buffer intrinsic noise, for example resulting from bursts of transcription of circuit components (Alon, 2007; Longo et al., 2013; Raser and O’Shea, 2005). However, data from flowering plants has shown that extrinsic fluctuations in auxin signaling input levels may be a significant source of noise in the auxin response GRN (Vernoux et al., 2011). To explore how the feedback loop between auxin signaling and tasiRNA-regulated ARFs affects the propagation of extrinsic auxin noise through the auxin response GRN in *Physcomitrella*, we adapted a computational model of the auxin response based on ordinary differential equations (ODEs) generated by Vernoux *et al*. (2011), and modified it to reflect the specific architecture of the moss auxin response GRN (**Figure 4B; Modeling Supplement SM1**). This model is based on a statistical mechanical approach to thermodynamic GRN modeling, as described by Bintu *et al.* (2005a), and is compatible with the timescales of development studied here. For a detailed discussion of the modeling approach and the parameters used, see the **Modeling Supplement**.

Two important measures of GRN function, susceptibility and noise amplification, were considered with and without transcriptional regulation of repressor ARFs by auxin signaling. Susceptibility (elsewhere also described as sensitivity or gain) measures the change in a system’s output relative to a small change in input (Bintu et al., 2005b; Hornung and Barkai, 2008). In this case, susceptibility reports the degree to which ARG expression changes in response to a sustained 1% change in auxin signal, and was assayed via numerical solutions of steady-state ARG transcript values.

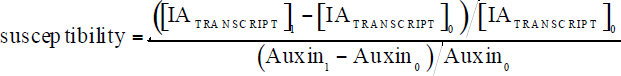

Noise amplification, on the other hand, measures the ratio between output noise and input noise (Hornung and Barkai, 2008). Here, it is indicative of the degree to which extrinsic auxin noise resulting from short fluctuations in auxin signaling levels are translated to fluctuations in ARG expression levels, and was assayed in computer simulations of the auxin response GRN.

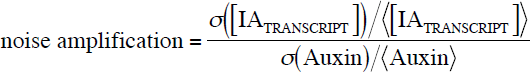

Feedback regulation from repressor ARFs onto the auxin response results in decreased susceptibility (**Modeling Supplement SM6A**), confirming predictions made in simpler genetic networks by both theoretical work (Paulsson, 2004) and simulations (Hornung and Barkai, 2008). However, there is no clear theoretical prediction regarding the effect of negative feedback on extrinsic noise amplification (Paulsson, 2004). Our model predicts that negative feedback from the *PpARFb* genes on the auxin response minimally affects extrinsic noise amplification. The small decrease in extrinsic auxin noise amplification that is observed is not significant (**Modeling Supplement SM6A**). Both findings are robust across a wide range of starting auxin input levels (**Modeling Supplement SM6C**). Considering the effect of multiple repressive feedback loops on intrinsic noise in other regulatory networks (Longo et al., 2013), this finding suggests that one advantage of this network architecture may be that it can buffer intrinsic noise without imparting strong amplification of extrinsic noise. Any benefits repressive feedback regulation in the auxin response GRN has in terms of promoting robustness against intrinsic fluctuations in gene expression are, however, coupled with a loss of susceptibility.

### tasiARF regulation promotes a robust auxin response

Small RNA regulation is likewise thought to lend robustness to the output of a GRN. Indeed, a recent study showed that small RNA regulation in mammalian cells suppresses intrinsic fluctuations in target gene expression (Schmiedel et al., 2015). However, the role of small RNA regulation in modulating extrinsic noise amplification is not known. To visualize how tasiRNA regulation affects the cellular response to auxin, we simulated the output of the auxin response network upon changes in auxin signal levels at two repressor ARF transcript degradation rate (γ_A-transcript_) values (see **Modeling Supplement**). The higher value, representing wild type, is based on the degradation rate of tasiRNA-targeted *ARF* transcripts in wild type *Arabidopsis* (Narsai et al., 2007). A 3.5-fold lower γ_A-transcript_ value, which results in a change in *PpARFb* level in line with that observed in *Ppsgs3*, is used to represent a lack of tasiARF regulation. Increasing γ_A-transcript_ results in a higher response output to a sustained step-increase in auxin signaling level (**Figure 4C**), reflecting increased susceptibility. In simulations of a noisy signal, increasing repressor ARF transcript degradation rate results in larger fluctuations in auxin-responsive gene expression (**Figure 4D**). This hints at an output of the auxin response that is more susceptible and less robust to extrinsic auxin fluctuations in the presence of tasiRNA regulation of PpARFb repressors.

However, while γ_A-transcript_ is higher in wild type than in *Ppsgs3*, precise ARF transcript degradation rates are not known in *Physcomitrella*. We thus modeled the effect of a spectrum of γ_A-transcript_ values on susceptibility and noise amplification. For a wide range of repressor *ARF* transcript degradation rates tested, increasing γ_A-transcript_ results in increased susceptibility to a small, sustained auxin signal that is correlated with an increase in extrinsic noise amplification in the auxin response GRN (**Figure 4E**). Substantiating these findings, modulation of ARF translation rates (π_A-_), which may also be affected by tasiRNA regulation, has the same linked effect on susceptibility and noise amplification levels (**Modeling Supplement SM5**). In addition, these outcomes are qualitatively robust to changes in all network parameter values (see **Modeling Supplement**). This generalizes the observation that tasiRNA regulation imparts susceptibility on the auxin response that is coupled with decreased robustness to auxin input noise. However, beyond a γ_A-transcript_ value of ~0.003, the relationship between these network properties change (**Figure 4E**). The auxin response GRN is too complex for an analysis of parameters contributing to this shift, but it is interesting to note that the best available estimates of network parameter values suggest that the wild type auxin response GRN may function close to this local maximum in susceptibility and extrinsic noise amplification (**Figure 4E**).

These findings further highlight the possible multifaceted contributions of small RNA regulation to GRN properties. While simulations predict that the small RNA-mediated regulation of PpARFb repressor ARFs impairs the plants’ ability to buffer ARG expression levels against fluctuations in auxin signaling input, small RNA regulation is expected to provide robustness against intrinsic noise in repressor ARF levels (Schmiedel et al., 2015). To experimentally assay the effect of tasiARF regulation on noise in the auxin response, we measured the coefficient of variation (CV) for transcript levels of six ARGs (Pires et al., 2013; Prigge et al., 2010) across biological replicates of wild type or *Ppsgs3*. Transcript levels for five of these genes, *PpIAA1a, PpIAA1b, PpIAA2, PpRSL5*, and *PpRSL6*, are more variable in *Ppsgs3* than in wild type (**Figure 4F**), and this effect is significant (P < 0.05) when measured across all six ARGs. This demonstrates that in *Physcomitrella*, the overall effect of tasiARFs on the auxin response GRN is to increase robustness of auxin-responsive gene expression, and seems to hint at different primary sources of noise in the *Arabidopsis* and *Physcomitrella* auxin response GRN.

### tasiARF regulation promotes phenotypic sensitivity to auxin

In the absence of exogenous auxin, *Ppsgs3* plants resemble auxin-resistant mutants in phenotype. However, this finding leaves open whether *Ppsgs3* is insensitive to auxin or responds with reduced sensitivity to changing auxin levels. The susceptibility effects suggest the latter. However, while susceptibility, which reports the change in response output resulting from a minute change in signaling input level (Bintu et al., 2005b; Hornung and Barkai, 2008), is a useful measure of network signaling properties, larger changes in input levels more accurately reflect auxin signaling *in vivo*. We therefore next modeled the qualitative effect of tasiRNA regulation on steady-state ARG expression across a broad range of increasing auxin signaling levels. As before, calculations were performed at two γ_A-transcript_ values, representing wild type (high γ_A-transcript_ and *Ppsgs3* (low γ_A-transcript_, respectively. For both values, the output of the simulated GRN to increasing auxin-signaling levels follows a sigmoid curve. However, the ARG transcript level is lower overall in the *Ppsgs3*-like regime, with differences between the two regimes especially pronounced at high auxin signaling levels (**Figure 5A**). The model thus predicts that tasiRNA regulation quantitatively sensitizes the auxin response across a wide range of auxin concentrations.

**Figure 5.**
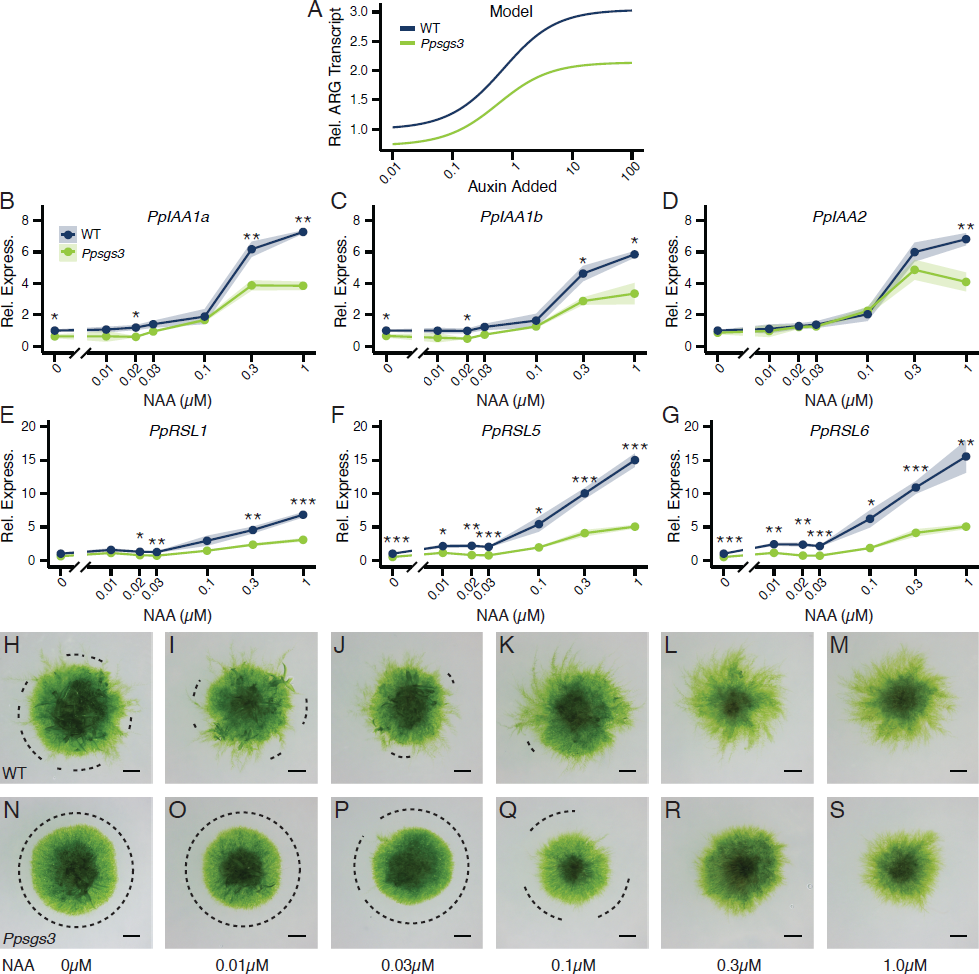
Perturbation of tasiRNA biogenesis dampens auxin sensitivity. (A) Model predictions for steady-state ARG transcript levels at increasing auxin concentrations for ‘wild type’ and *’Ppsgs3\* normalized to ‘wild type’ levels at baseline auxin signaling input. X-axis, value of auxin signaling input added to the baseline level in the model (see **Modeling Supplement**). (B-G) Relative transcript levels (mean ± SE, n ≥3), normalized to wild type grown on media without added NAA, for *PpIAAla* (B), *PpIAAlb* (C), *PpIAA2* (D), *PpRSLl* (E), *PpRSL5* (F), and *PpRSL6* (G) in 15 day-old wild type and *Ppsgs3* plants grown on increasing NAA concentrations. *P < 0.05, **P < 0.01, ***P < 0.001, Student’s *t* test. (H-S) Caulonema occurrence on15 day-old wild type (H-M) and *Ppsgs3* (N-S) plants grown on media supplemented with increasing concentrations of NAA. Dotted lines, protonemal regions lacking caulonema. Scalebar, 1 mm.

To experimentally test this model prediction, we measured the transcript level profiles for *PpIAA1a, PpIAA1b, PpIAA2, PpRSL1, PpRSL5*, and *PpRSL6* in wild type and *Ppsgs3* plants grown on media supplemented with between 0 - 1μM NAA (**Figure 5B-G**). Very low doses of exogenous auxin not commonly assayed were included in this range to observe the effect of subtle changes in auxin levels on ARG expression. As expected, expression of all six genes in wild type increases upon treatment with NAA. Consistent with computational predictions, the *PpIAA* gene expression profiles follow a sigmoidal curve, with the steepest increase in auxin-induced expression occurring between 0.1 and 0.3μM NAA. The change in expression for the *PpRSL* genes appears more gradual, perhaps reflecting the complex transcriptional interaction between RSL family members (Pires et al., 2013). Importantly, *Ppsgs3* plants display reduced sensitivity to auxin but, as predicted by the model, still demonstrate a clear auxin response. Transcript levels for all six ARGs are significantly upregulated in response to treatment with auxin concentrations greater than 0.1μM NAA, but overall remain lower than seen in wild type. Significant transcript level differences between wild type and *Ppsgs3* can be observed even at low auxin levels, despite increased expression variability in *Ppsgs3*.

The decreased sensitivity of *Ppsgs3* plants to auxin at the molecular level is reflected at the phenotypic level. As described above, 15 day-old wild type plants grown without exogenous auxin produce multiple caulonemal filaments around their circumference. The number of these filaments increases visibly in wild type plants grown on 0.01- 0.1μM NAA, with caulonema emerging along most of the protonemal circumference upon treatment with 0.1μM NAA (**Figure 5I-K**). On auxin concentrations beyond that, the protonemal network is increasingly comprised of caulonema, resulting in a sparser and lighter appearance (**Figure 5L, M**). In contrast, *Ppsgs3* plants grown on auxin concentrations up to 0.1 μM NAA show a minimal induction of caulonemal differentiation, and continue to form dense chloronemal networks with few or no caulonemal filaments (**Figure 5O, P**). Treatment with levels of NAA over 0.1 μM, however, can override this defect, such that caulonemal filaments differentiate along the entire protonemal circumference (**Figure 5R, S**). These findings reaffirm that *Ppsgs3* plants are not generally impaired in their ability to form caulonemal filaments or respond to exogenous auxin. Rather, *Ppsgs3* mutants display a reduced auxin sensitivity such that a smaller proportion of chloronemal cells differentiates into caulonema in response to a given auxin signal.

These phenotypic and gene expression data parallel the computational predictions of single-cell auxin responses and demonstrate that tasiRNAs play a key role in sensitizing the auxin response in *Physcomitrella* via their regulation of *PpARFb* genes. By spatially regulating *PpARFb* expression, tasiARFs, in concert with miR1219, create a zone of variable auxin sensitivity at the protonemal edge, allowing the ratio of caulonemal to chloronemal cells to be tuned in response to increasing auxin levels.

### *Ppsgs3* plants have reduced sensitivity to environmental cues that guide development

Quantitative regulation of development, such as that observed above in the *Physcomitrella* auxin response, is especially important in plants, which due to their sessile nature must integrate a wide range of environmental cues into their developmental programs. Caulonemal differentiation appears to be especially labile to environmental regulation and is, for instance, modulated by light quality and levels of available nutrients (Reski, 1998). We therefore hypothesized that, as quantitative regulators of caulonemal differentiation, tasiARFs may be important for developmental plasticity and the ability to adjust development in response to environmental change. To test this possibility, we analyzed the effect that reduced substratum nitrogen has on wild type and *Ppsgs3* protonemal network morphology. For wild type, decreasing nitrogen levels in the growth medium from the typical 5 mM di-Ammonium Tartrate (dAT) promotes the chloronemal-to-caulonemal transition, such that the proportion of caulonemal filaments on plants grown on media containing 1, 0.2, 0.05, or 0 mM dAT increases substantially (**Figure 6A-D**). Although the number of caulonemal filaments on *Ppsgs3* plants also increases with lower nitrogen levels in the media, *Ppsgs3* plants grown on 0.2 mM dAT and especially 1 mM dAT develop proportionally fewer caulonema than wild type (**Figure 6E-H**). A diminished responsiveness to changes in substratum nitrogen levels is also seen in plants expressing the small RNA-insensitive *PpARFb4-GUS-m*t\** variant (**Figure 6I-L**), pointing to a role for tasiARF specifically in modulating the plant’s ability to tune development in response to this key environmental cue.

**Figure 6.**
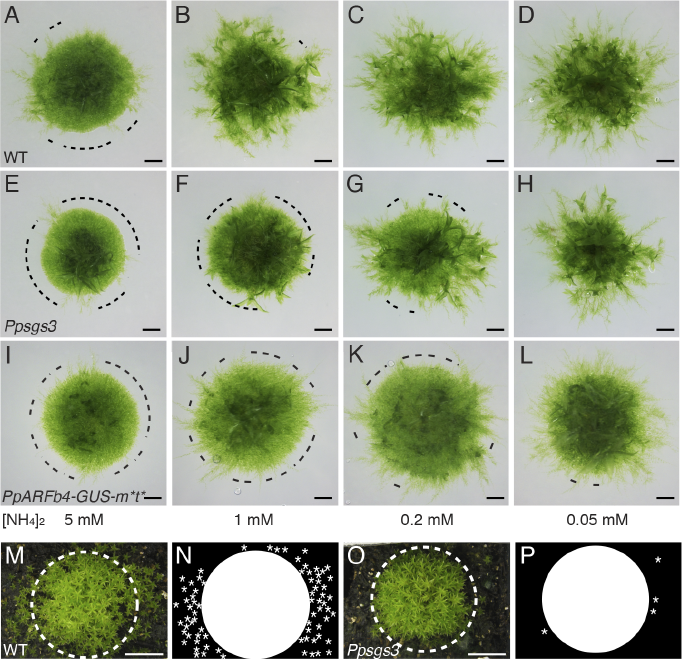
*Ppsgs3* has a decreased sensitivity to changes in substratum nitrogen levels. (A-J) Caulonema occurrence on 15 day-old wild type (A-D), *Ppsgs3* (E-H), and *PpARFb4-GUS-m*t\** (I-L) plants grown on BCD media supplemented with decreasing concentrations of diAmmonium Tartrate (dAT). Dotted line, protonemal regions lacking caulonema. Scalebar, 1mm. (M-P) Gametophore distribution on 2 month-old soil-grown wild type (M, N) and *Ppsgs3* (O, P) plants. Dotted line, approximate edge of the main wild type plant, represented by the circle in N and P; asterisks, positions of gametophores away from the main plant’s edge. Scalebar, 1cm.

*Physcomitrella*’s natural habitat is, however, vastly different from the agar plates on which moss is grown in the lab, and includes agricultural fields and the moist soil at the edges of bodies of freshwater. To test how tasiRNA regulation impacts moss development in conditions more reflective of *Physcomitrella*’s natural environment, we compared the growth of wild type and *Ppsgs3* in moist soil. While it was not possible to directly observe protonemal development, two months after transplantation to soil, wild type plants reveal a tight central cluster of gametophores, ~2 cm wide, surrounded by numerous dispersed gametophores (**Figure 6M, N**). By contrast, *Ppsgs3* plants have a slightly smaller central gametophore cluster and few, if any, dispersed gametophores (**Figure 6O, P**). This result suggests that the decrease in caulonemal differentiation caused by the loss of tasiRNA biogenesis severely impacts the ability of the plant to colonize its substratum, highlighting the importance of tasiRNAs in moss’ interactions with its environment.

Taken together, our work shows that the *TAS3* tasiRNA pathway spatially restricts expression of a conserved class of repressor ARF targets to a variable subset of cells at the protonemal edge. tasiARF-mediated ARFb regulation alters the signaling properties of the auxin response GRN, conferring sensitivity and robustness onto the auxin response. The tasiRNA pathway thus tunes the auxin sensitivity of cells at the edge, leading to a stochastic pattern of caulonemal filament differentiation and a variable developmental response to changing environmental signals.

## DISCUSSION

### An ancestral role for *TAS3* tasiRNAs in modulating auxin-regulated processes via ARFb repressor proteins

Instances of repeated cooption of specific genetic networks to direct new developmental processes are common across evolution (Carroll et al., 2004a; Plavskin and Timmermans, 2012). This study considers network properties that promote recurrent cooption by investigating one often-repurposed genetic pathway, the *TAS3* tasiRNA pathway, and its targets. Although this small RNA pathway likely originated in the common ancestor of all land plants (Axtell et al., 2007), its known roles in flowering plants are in a diverse set of recently evolved processes, including flower, root, and leaf development (see Plavskin and Timmermans, 2012). Our results demonstrate that in the moss *Physcomitrella patens*, the *TAS3* tasiRNA pathway acts in the gametophytic stages of development to modulate gametophore initiation, protonemal branch determinacy, and caulonemal differentiation. The roles of tasiRNAs in protonemal development likely represent an independent cooption event, rather than an ancestral function, as the extensive and complex protonemal network found in *Physcomitrella* evolved within the moss lineage (Mishler and Churchill, 1984). Nonetheless, the functional targets of the *Physcomitrella* tasiRNA pathway are conserved. Its effects on gametophyte development are via regulation of B-family repressor ARFs, whereas the functions of novel tasiRNA targets, such as the AP2- and ZF-family transcription factors (Arif et al., 2012; Axtell et al., 2007; Talmor-Neiman et al., 2006), remain unknown and are not immediately apparent from the *Ppsgs3* phenotype. Importantly, tasiRNAs in flowering plants likewise affect development via regulation of members of the B-group of ARF genes (Chitwood et al., 2009; Dotto et al., 2014; Hunter et al., 2006; Yifhar et al., 2012; Zhou et al., 2013). Transcription factors in this clade are known to modulate the auxin response in space and time by repressing ARG expression (Finet and Jaillais, 2012; Vernoux et al., 2011). Our findings thus point to an ancestral role of the *TAS3* tasiRNA pathway in regulating ARFb expression, with the tasiARF-ARFb module repeatedly coopted over the course of plant evolution to modulate the auxin response GRN in select and diverse developmental contexts.

### tasiARFs promote sensitivity and robustness of the auxin response

The strong temporal fluctuations in auxin signaling levels observed in flowering plant meristems suggested that the need to buffer this extrinsic noise may be a key factor shaping the organization and evolution of the auxin response GRN (Vernoux et al., 2011). Our findings predict other properties may have driven the tasiARF-ARFb module’s repeated cooption. Our model predicts that negative feedback from repressor *ARFb* genes onto the auxin response, which is at least in part conserved in flowering plants (Marin et al., 2010; Yoon et al., 2010), minimally affects extrinsic noise amplification. Instead, a predicted advantage of this network feature is that it may limit intrinsic noise resulting from the inherent variability in gene expression (Alon, 2007; Longo et al., 2013), although this prediction is based on models of different GRNs. A key contribution of the *TAS3* tasiRNA pathway to the auxin response GRN during *Physcomitrella* protonemal development may also be to confer robustness onto the auxin response, as loss of tasiRNA regulation was found to increase variation in ARG expression. However, this contribution seems to reflect a suppression of intrinsic rather than extrinsic noise in the auxin response GRN, as our model predicts that tasiRNA regulation amplifies the extrinsic noise from fluctuating auxin signaling input levels. On the other hand, a role for small RNAs in repressing intrinsic noise is in line with recent findings in mammalian cells (Schmiedel et al., 2015). Interestingly, Schmiedel *et al*. (2015) demonstrated that intrinsic noise repression by small RNAs is especially effective when transcripts contain multiple small RNA binding sites, as is the case for the tasiRNA-targeted *ARFb* genes in moss as well as flowering plants (Chitwood et al., 2009; Dotto et al., 2014; Yifhar et al., 2012; Zhou et al., 2013).

Although tasiRNAs cannot affect development without their targets, repressor ARFs can, and do, function in the absence of tasiRNA regulation (Plavskin and Timmermans, 2012). The repeated repurposing of the tasiARF-ARFb module over the course of plant evolution therefore suggests that introducing this module into a novel, auxin-regulated context may provide a selective advantage. Both the increased sensitivity and robustness lent to the auxin response by the tasiRNA pathway promote the faithful transfer of information through a signaling network. Evidence that selection may indeed act on such network properties has been identified across the tree of life, and includes examples of selection acting on regulatory mutations that minimize expression noise in yeast (Metzger et al., 2015), as well as the conservation of shadow enhancers that maintain robust gene expression in animal systems (Frankel et al., 2010; Hong et al., 2008). The network properties lent to the auxin response by the tasiARF-ARFb module uncovered in this work thus provide a compelling explanation for the repeated cooption of this module throughout land plant evolution. Furthermore, these signaling properties may provide an explanation for the prevalence of evolutionarily conserved small RNA-target modules in plants as well as animals.

Considering these points, it is interesting to note that a subset of the flowering plant B-group *ARF* genes have lost tasiRNA regulation (Plavskin and Timmermans, 2012). Our modeling results provide a possible explanation for this diversification. As plants evolved the need to regulate auxin responses in organs where fluctuations in auxin levels are high, such as in meristems (Vernoux et al., 2011), the benefit provided by tasiRNAs in dampening intrinsic expression noise may have been offset by their amplification of extrinsic auxin input noise. Likewise, despite the apparent importance of small RNA regulation in promoting sensitivity and robustness of the auxin response, many auxin-regulated processes in *Physcomitrella* development, such as in the leafy gametophore and sporophyte (Bennett et al., 2014; Viaene et al., 2014), are unaffected when tasiRNA function is perturbed. An interesting direction for future studies may be to explore whether miR160 and its highly conserved targets in the C-family of repressor ARFs (Axtell and Bartel, 2005; Finet et al., 2013; Plavskin and Timmermans, 2012), act in place of the tasiARF-ARFb module in the regulation of these processes.

### tasiARF-mediated PpARFb regulation allows for stochastic protonemal cell fate determination

In addition to the network properties acting at the cellular level, the tasiARF-mediated regulation of PpARFb expression generates stochasticity at the whole plant level. Moss protonema is a heterogenous tissue, with caulonemal filaments specified at seemingly random locations along the edge of the protonemal network over the course of development. Well-known examples of tissues where cell fates are specified using a stochastic choice mechanism to create a randomly mixed population of cell types exist in the metazoan nervous system, and include the diversification of olfactory and visual sensory neurons (Johnston and Desplan, 2010). However, the source of stochasticity in these examples is unclear.

Our observations indicate that tasiARF, in conjunction with miR1219, regulates caulonemal fate specification in *Physcomitrella* by limiting PpARFb expression to a random subset of chloronemal tip cells. To create variability in PpARFb expression across individual filament tips, the levels of small RNAs in the protonema must be carefully tuned. In this regard it may be interesting that in *Arabidopsis*, the cell-to-cell movement of tasiARF from a localized source creates a small RNA gradient across developing leaf primordia (Chitwood et al., 2009). A gradient of tasiARF and miR1219 activity that dissipates towards the filament tips, perhaps generated by small RNA movement from a source at the center of the protonemal network, presents a possible mechanism to generate stochastiocity in *PpARFb* expression at the protonemal edge. The fact that the number of cells expressing PpARFb varies from filament to filament can also be explained by local subtle variation in the shape of a small RNA activity gradient. If so, patterning by small RNA gradients may represent a novel mechanism of stochastic cell fate specification.

Stochastic cell fate decisions are often important for environmental plasticity, allowing ‘bet-hedging’ of cell fate choices within populations of cells (Johnston and Desplan, 2010). In combination with the network properties small RNA regulation confers on the auxin response GRN, stochasticity in caulonema specification resulting from spatial variation in tasiARF and miR1219 regulation of PpARFb proteins appears important for the developmental response to environmental cues. To fine-tune morphology to subtle gradations in environmental conditions, organisms must modulate development in a quantitative, graded manner. However, to create such gradation in a binary system, such as the chloronemal-caulonemal cell fate switch, cells must display variability in their response to the switch-inducing signal (Ferrell and Machleder, 1998). By limiting PpARFb expression to a random subset of chloronemal tip cells, the tasiRNA pathway, along with miR1219, creates variable auxin responsiveness at the edge of the developing protonema. Tip cells with comparatively high small RNA activity and low repressor ARFb levels have increased auxin sensitivity, and are thus more likely to differentiate into caulonema in response to auxin signaling. Our finding that the number of caulonemal filaments increases progressively with increasing auxin levels supports this notion. The decreased response to small changes in substratum nitrogen availability in moss plants defective in tasiRNA biogenesis further underscores the importance of tasiRNAs in maintaining a plastic response to the environment.

With the role of tasiRNAs in sensitizing development to environmental inputs in mind, parallels can be drawn between independently evolved tasiRNA-regulated processes in mosses and flowering plants. Lateral root initiation in *Arabidopsis*, although likely sharing little with protonemal development in terms of cellular mechanisms, is also auxin- and tasiRNA-regulated, and is carefully tuned by environmental inputs, including substratum nitrogen levels (Gifford et al., 2008). Likewise, the auxin- and tasiRNA-regulated specification of abaxial-adaxial polarity in flowering plant leaves is sensitive to environmental inputs, with the number of adaxial and abaxial cell layers regulated in part by light quality (Kozuka et al., 2011).

### Network signaling properties as drivers of GRN cooption

Although the mechanisms by which cooption of genetic networks occurs have been extensively investigated, the reasons for the preferential repeated cooption of select networks are less well understood. One potential benefit of GRN cooption is that it allows the redeployment of a ‘differentiation gene battery’ involved in a specific cellular or developmental process (Erwin and Davidson, 2009). For example, the *RSL* genes were repeatedly repurposed during the evolution of plant organs that develop via filamentous growth, such as protonema in moss and root hairs in flowering plants (Jang et al., 2011; Menand et al., 2007; Pires et al., 2013). In contrast, we propose that the repeated cooption of the *TAS3* tasiRNA pathway, together with its ARFb targets, was driven by the properties that small RNA regulation lends to the auxin response, rather than by the redeployment of a specific downstream developmental process. We find that tasiRNAs lend the networks they regulate two key properties: robustness to intrinsic noise in target levels, and sensitivity to environmental signals. Furthermore, small RNAs may provide a mechanism to create stochastic cell fate patterns that are developmentally plastic and increase the spectrum of responses to environmental stimuli. Considering the signaling properties of a genetic network, and not just its developmental output, may thus be critical to understanding the evolution of complex multicellular forms.

## MATERIALS AND METHODS

The transformation and propagation of *Physcomitrella* was performed as described (Cove et al., 2009), with minor adjustments detailed in Supplementary Materials and Methods. For phenotypic or gene expression level measurements, protonema were subcultured on cellophane plates 2-5 times, and plantlets 1-3 mm in diameter transplanted to BCDAT plates at pH 5.8 containing 0.9 μM FeSO4. For chemical treatments, auxin, PCIB, or estradiol was added at the indicated concentrations. Nitrogen level experiments were performed on BCD plates supplemented with the appropriate amount of diammonium tartrate. For soil experiments, plants were grown on Rediearth (Sungro) under a plastic dome for two months at 22°C under a 16/8 light/dark regime. Cell size measurements were performed on chloronemal cells 3-5 cells from filament tips. Branching was analyzed on 3 week-old plants. For gametophore counts, the number of gametophore buds with at least 1 phyllid were counted. Histochemical staining was performed on 15 day-old plants as described (Chitwood et al., 2009). For all qRT-PCR and 5’ RACE experiments, total RNA was extracted using Trizol reagent (Invitrogen) according to the manufacturer’s suggested protocol. Importantly, unless otherwise stated, RNA was extracted from whole 15 day-old plants as detailed in Supplementary Materials and Methods. Small RNA qRT-PCR was performed following a protocol modified from Varkonyi-Gasic et al. (2007), and RLM 5’ RACE was performed as described in Axtell et al. (2007). All expression levels were calculated relative to untreated wild type controls, after normalization to levels of GAPDH (gene expression) or U6 (small RNA expression). Statistical significance was calculated from at least three biological and two technical replicates using student’s t-test between wild type and mutant samples within a given condition. For detailed experimental procedures, see Supplemental Experimental Procedures. For detailed computational modeling procedures, see Supplemental Modeling Information.

## ACKNOWLEDGEMENTS

We thank members of the Timmermans, Quatrano, and Hasebe laboratories who have contributed ideas and thoughtful comments to the manuscript, Tim Mulligan for plant care, and Elijah Salome-Diaz, Carol Hu, Cristina Marco, and members of the Hasebe lab for technical assistance. We also thank Teva Vernoux for sharing modeling code. YP was funded by an Al Hershey fellowship from the Watson School of Biological Sciences and a predoctoral training grant 5T32GM065094 from the NIGMS. This work was also supported by the EAPSI grant OISE-1015679 from the NSF and the JSPS. MH is funded by ERATO and KAKENHI from MEXT, Japan. Research on small RNA regulation in the Timmermans lab is supported by grants from the NSF (IOS-1355018 and MCB-1159098) and an Alexander von Humboldt Professorship from the DFG.

### AUTHOR CONTRIBUTIONS

YP and MT conceived the study and designed the project. YP and MT planned the experiments with input from P-F.P, AN, MH, and RQ. YP, P-F.P, and AN generated the transgenic moss strains. YP performed the experiments. YP performed the computational modeling with input from GA and MT. YP and MT wrote the manuscript.

